# Systematic *in vivo* quantification of microRNA affinities

**DOI:** 10.1101/2021.03.19.436154

**Authors:** Sebastian Brosig, Hanna L. Sladitschek, Pierre A. Neveu

## Abstract

The majority of mammalian genes are under regulation by microRNAs, yet predicting the extent of miRNA-mediated repression has remained elusive. Here we systematically quantified the biological impact of miRNAs conserved in vertebrates using stable mouse embryonic stem cell lines expressing sensitive fluorescent reporters. Differentiation of these 163 lines to the three germ layers revealed that the majority of conserved miRNAs have detectable changes in activity. We determined *in vivo* target affinity K_D_ of 115 miRNAs by integrating activity measurements, CRISPR/Cas miRNA knockouts and miRNA sequencing. Target affinities of individual miRNAs spanned several orders of magnitude, with highly expressed miRNAs having overall higher K_D_. Scaling miRNA expression levels by their respective K_D_ recapitulated the relative number of Argonaute-bound targets for individual miRNA families. Our results provide a rationale to determine the set of miRNAs with a biological activity in a given cell type, K_D_ values setting expression thresholds for target repression.

## Introduction

MicroRNAs (miRNAs) are a family of small non-coding RNAs that post-transcriptionally regulate gene expression (Bartel, 2009). miRNAs are incorporated in the Argonaute-containing RNA-induced silencing complex (Fabian *et al*, 2010) and direct RISC to target mRNAs predominantly in their 3’-untranslated regions (3’-UTRs) (Bartel, 2004). Base pairing of the nucleotides 2-7 at the 5’-end of the miRNA named the “seed” region to the 3’-UTR of mRNAs is generally sufficient for miRNA target recognition, which results in either translational repression of the target (Carthew and Sontheimer, 2009) or target mRNA degradation (Guo *et al*, 2010).

There is an evolutionary conservation of miRNA families (Grimson *et al*, 2008; Griffiths-Jones *et al*, 2008) and their corresponding targets (Friedman *et al*, 2009). For example, both the let-7 sequence and its temporal expression during development are highly conserved across bilaterians (Pasquinelli *et al*, 2000). Moreover, let-7 was shown to be a pro-differentiation regulator from *C. elegans* (Reinhart *et al*, 2000) to mouse (Viswanathan *et al*, 2008).

Most mRNAs are regulated by miRNAs (Lewis *et al*, 2005), with individual miRNAs targeting hundreds of different transcripts (Grimson *et al*, 2007; Lim *et al*, 2005). mRNAs that are actually regulated by miRNAs in a cell can be identified by characterizing the Ago-bound RNAs (Hafner *et al*, 2010; Chi *et al*, 2009). Such datasets enable the prediction of binding affinities of a specific miRNA for different target sites (Khorshid *et al*, 2013). It is also well established that an increase in miRNA seed base pairing results in higher binding affinity (Nielsen *et al*, 2007; Friedman *et al*, 2009). However, the differences in binding affinities of individual miRNAs are poorly characterized. *In vitro* experiments using miRNA-loaded complexes determined that individual miRNAs have different affinities to a model target and that the binding properties are uniquely influenced by Ago2 (Chandradoss *et al*, 2015; Salomon *et al*, 2015; Wee *et al*, 2012; McGeary *et al*, 2019). Despite the conservation of both miRNA families and their targets, large scale studies aiming at assessing miRNA impact on synthetic targets concluded that the majority of miRNAs had no measurable activity (Mullokandov *et al*, 2012; Gam *et al*, 2018). Here we generated stable mouse embryonic stem cell (mESC) lines expressing sensitive fluorescent reporters (Sladitschek and Neveu, 2016) for the 163 miRNAs conserved in vertebrates. Systematic differentiation of these lines to the three germ layers, mesoderm, endoderm and ectoderm, revealed that the majority of conserved miRNAs have measurable changes in biological activity. Furthermore, the combination of miRNA activity measurements, CRISPR/Cas miRNA knockouts with miRNA deep sequencing allowed us to determine the *in vivo* binding affinity K_D_ of the miRNA:target interaction for 115 miRNAs. We found that K_D_ for individual miRNAs spanned several orders of magnitude and that K_D_ was weakly correlated to the median miRNA expression levels in mouse tissues. In order to establish the relevance of our measurements for endogenous targets, we analyzed published Ago-CLIP data (Bosson *et al*, 2014). A mathematical model in which miRNA expression levels were normalized by their respective K_D_ accounted for the abundance of the endogenous Ago-bound target pool. Therefore, miRNAs have to be expressed at levels higher than their respective K_D_ to elicit a significant impact on target gene expression.

## Results and Discussion

In order to monitor miRNA activity in a variety of cell types, we relied on a fluorescence-based ratiometric miRNA activity sensor, with stable expression during mESC differentiation (Sladitschek and Neveu, 2016). This reporter is designed around a bidirectional CAG promoter driving the expression of two genes encoding for two fluorescent proteins: one gene contained in its 3’-UTR a binding site for the miRNA to be monitored and served as detector channel, while the other gene served as normalizer channel to correct for variation in reporter expression (Fig 1A). Binding of the miRNA to the detector transcript will lead to a repression of its expression (Fig 1A). Such a reporter construct can be stably integrated in the genome of mESCs (Fig 1B). Computing the reporter ratio as the intensity in the detector channel divided by the intensity in the normalizer channel in a single cell is a measure of the activity of a specific miRNA in that particular cell. Changes in miRNA levels can then be read out by a change in detector signal of opposite sign (Fig 1C). In the case of reporters relying on a single binding site in the 3’-UTR of the detector mRNA, we previously determined (Sladitschek and Neveu, 2015) that the reporter ratio depended with a Hill function 1/1+M/K_D_ on the miRNA levels M and the affinity K_D_ for the reporter construct (Fig 1D). The absence of cooperativity leads to a more graded response compared to constructs employing several binding sites, which would make the reporter response ultrasensitive around K_D_. In particular, K_D_ can be determined by measuring the reporter response while the levels of the assessed miRNA are changed experimentally (Fig 1D).

**Figure 1.**
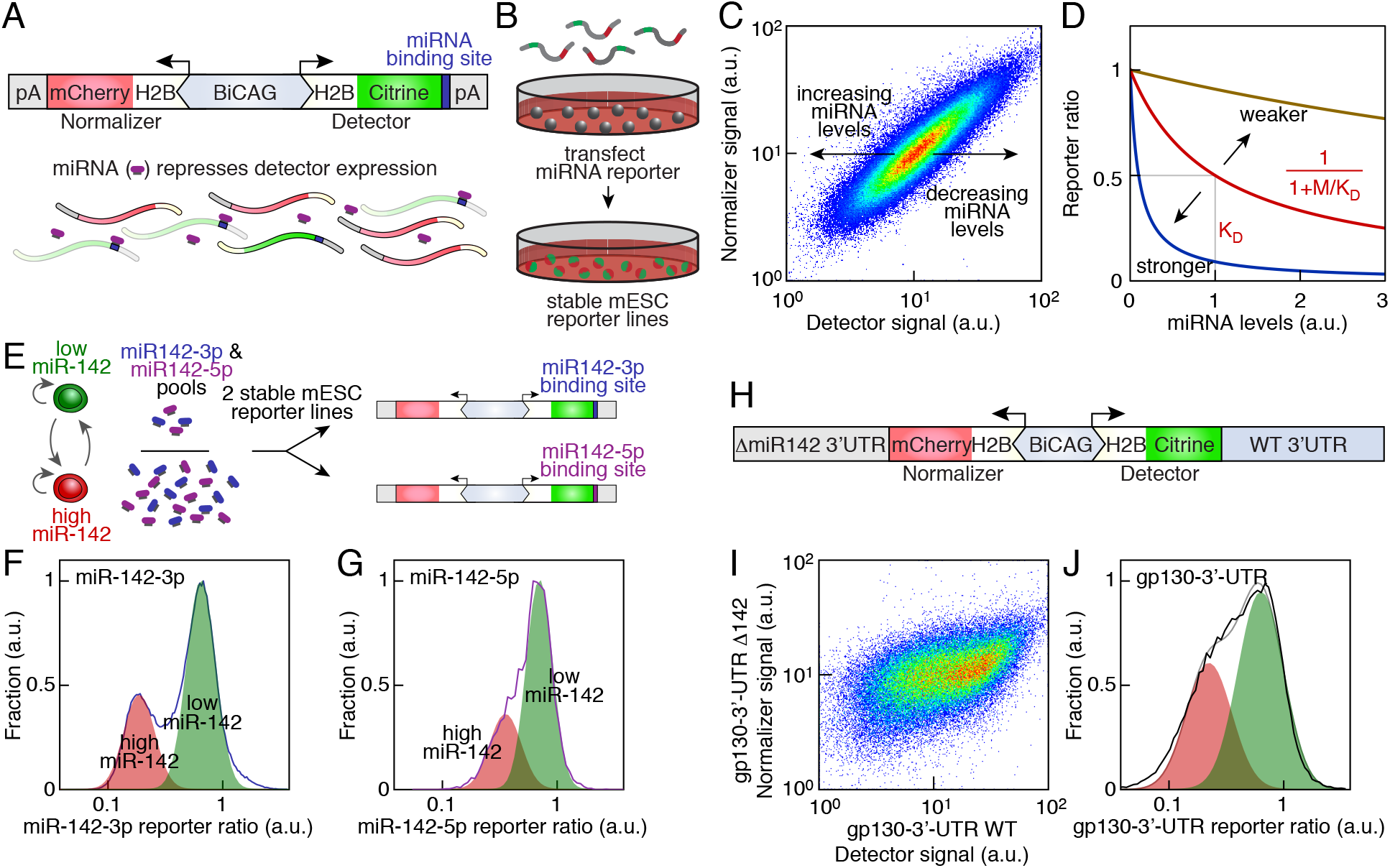
miRNAs have different affinities for a model target *in vivo*. A Scheme of the single-cell fluorescent reporter of miRNA activity. B Stable mESC reporter lines were established for all 163 miRNAs conserved in vertebrates. C Detector and normalizer signals as measured by flow cytometry in an mESC reporter line stably expressing a miR-136-3p reporter. D Scheme of the dependence of the reporter ratio on miRNA levels (M) and target affinity (K_D_). E miR-142 is bimodally expressed in self-renewing mESCs leading to two subpopulations with different sizes of the miR-142-3p and miR-142-5p pool. These can be assessed with miR-142-3p or miR-142-5p reporter lines. F Distribution of the miR-142-3p reporter ratio in self-renewing mESCs. The experimental data was approximated with two log-normal distributions corresponding to high and low miR-142 mESCs. G Distribution of the miR-142-5p reporter ratio in self-renewing mESCs. The experimental data was approximated with two log-normal distributions corresponding to high and low miR-142 mESCs. H Scheme of the reporter to measure the contribution of a specific miRNA to the regulation of a 3’-UTR. The normalizer gene is fused to the 3’-UTR with deletion of the binding sites of the considered miRNA while the detector is fused to the wild type 3’-UTR. I Detector and normalizer expression in a clonal mESC population stably expressing a gp130 3’-UTR reporter. The detector is fused to a wild type gp130 3’-UTR while the normalizer is fused to a gp130 3’-UTR with deleted miR-142 binding sites (Δ142). J Distribution of a gp130 3’-UTR reporter ratio in self-renewing mESCs. The experimental data was approximated with two log-normal distributions (black line: experimental data, gray line: fit with two log-normal distributions).

We first determined how different miRNA regions impacted the reporter expression. Taking miR-136-5p as a model miRNA, we found that a single binding site containing only the miR-136-5p seed region elicited the same repression of the reporter as a single binding site fully complementary to the entire miR-136-5p (Fig S1A and S1B), which is in line with *in vitro* binding assays using mouse AGO2 (Wee *et al*, 2012). A binding site complementary to the 3’ region of miR-136-5p (lacking the seed region) had however no effect on the reporter expression (Fig S1C). These results were consistent with Ago2 exposing the miRNA seed region for binding to an mRNA target (Elkayam *et al*, 2012; Nakanishi *et al*, 2012; Schirle and MacRae, 2012). We thus retained a single binding site fully complementary to the miRNA which activity is to be measured. This was shown to trigger detector mRNA degradation (Mullokandov *et al*, 2012), preventing sponging out the miRNA (Ebert *et al*, 2007; Mukherji *et al*, 2011). Weakly expressed miRNAs would be most sensitive to titration effects. Using miR-142-3p as a model (with expression levels in the range of 100s reads per million), we found that the reporter responded with less than 2.3% variation over a 20-fold expression range (Fig S1D). The miRNA reporter was therefore not influenced by any titration effects from the reporter construct itself.

We took advantage of the known bimodal expression of miR-142 in mESCs under self-renewing conditions (Sladitschek and Neveu, 2015) to generate well characterized variations in miRNA expression levels that can be assessed in the same sample. Indeed two populations of cells coexist: one with high levels of miR-142, and the other one with low levels of miR-142 (Fig 1E). The two mature forms derived from the pre-miR-142 stem loop miR-142-3p and miR-142-5p had highly correlated expression levels (Fig S1E) and therefore the same difference in expression between the two mESC subpopulations (Sladitschek and Neveu, 2015). We therefore compared miRNA activity in reporter lines for miR-142-3p or miR-142-5p (Fig 1E). While the distribution of the log-transformed reporter ratio was well approximated by the sum of two normal distributions in both cases, the two means were more separated for the miR-142-3p reporter compared to the miR-142-5p reporter (Fig 1F and 1G). Using the miRNA reporter response curve (Fig 1D), we determined that miR-142-3p had 1.5 times smaller target affinity than miR-142-5p. Thus miRNAs had different target affinities *in vivo*.

In order to show that miRNAs have similar effects on endogenous targets, we created a reporter line for the 3’-UTR of gp130, a known target of miR-142 (Sladitschek and Neveu, 2015). The detector color was fused with a wild type gp130 3’-UTR while the normalizer color had a gp130 3’-UTR with mutated miR-142 binding sites (Fig 1H). At a given normalizer level, the detector signal in single cells spanned a range larger than 10-fold in response to the intrinsic variability in miR-142 expression levels in mESCs under self-renewing conditions (Fig 1I). This regulation was mostly at the post transcriptional level as gp130 mRNA levels varied by less than 20% between high and low miR-142 cells (Fig S1F). Like for miR-142-3p and miR-142-5p, the distribution of the log-transformed ratio of the gp130 3’-UTR reporter was well approximated by two normal distributions (Fig 1J). Moreover, the repression span due to the action of miR-142 on gp130 3’-UTR was similar to the one observed with our synthetic miR-142-3p activity reporter. Thus miRNAs can elicit large effects on the expression of endogenous targets.

We went on to assess if we could measure changes in miRNA activity *in vivo*, taking advantage of the different miRNA expression profiles between pluripotent cells and differentiated cells (Neveu *et al*, 2010). The downregulation of pluripotency-specific miRNAs during mESC differentiation was used as a benchmark. We differentiated to mesoderm an mESC line reporting on the activity of the miRNA miR-295-3p, which is expressed exclusively in mESCs. Compared to undifferentiated mESCs, there was a large shift in miR-295-3p detector signal at the end of the differentiation procedure (Fig S2A and S2B). In fact, we could detect a progressive increase in the miR-295-3p reporter ratio while mESCs adopted mesoderm, endoderm or ectoderm fates (Fig S2C–E). It demonstrated our ability to measure changes in miRNA activity during differentiation.

Our standardized miRNA reporter removes the potential accessibility problems associated with binding sites in endogenous targets because the only variable part in the construct is the miRNA binding site, the flanking regions remaining the same. It therefore enables to compare the activity of different miRNAs in a similar context. We thus generated reporter mESC lines for each of the 163 miRNAs found in the mouse genome that have seeds conserved across vertebrates (Grimson *et al*, 2007). Sequencing the miRNA complement during germ layer acquisition established that the majority of conserved miRNAs changed their expression levels during differentiation (Fig S3A and S3B). For the biologically active miRNAs, these expression changes should be accompanied by changes in miRNA activity. The 163 mESC reporter lines were then systematically differentiated to the three germ layers (Fig 2A). Normalizer and detector signals were measured by flow cytometry in single cells every day while the cells were undergoing differentiation (Fig 2A). The reporter ratio was then computed and its mean served as an approximation of the activity of the assessed miRNA in the entire population (Fig 2B). Surprisingly, the vast majority of miRNAs displayed a change in reporter ratio during germ layer fate acquisition (Fig 2C). Using principal component analysis on the dataset of reporter ratios, we found that the specification to the different germ layers followed their own trajectory in the space of miRNA activity (Fig S3C). This reflected the individual trajectories that diverged from the onset of the differentiation procedure in the expression space of both miRNAs (Fig S3D) and protein-coding genes (Sladitschek and Neveu, 2019). Having conducted both miRNA-Seq and mRNA-Seq from the same isolated total RNA, we analyzed the genome-wide effects of miRNA expression on the cell’s transcriptome. There was an anti-correlation between the expression levels of a given miRNA and of target mRNAs possessing more than four predicted binding sites for this specific miRNA (Fig S3E). Interestingly, no such anti-correlation was observed for mRNAs with a single predicted target site. Thus, the widespread changes in miRNA activity were accompanied by a genome-wide impact of miRNA expression in the relative downregulation of the mRNA levels of their mRNA targets.

**Figure 2.**
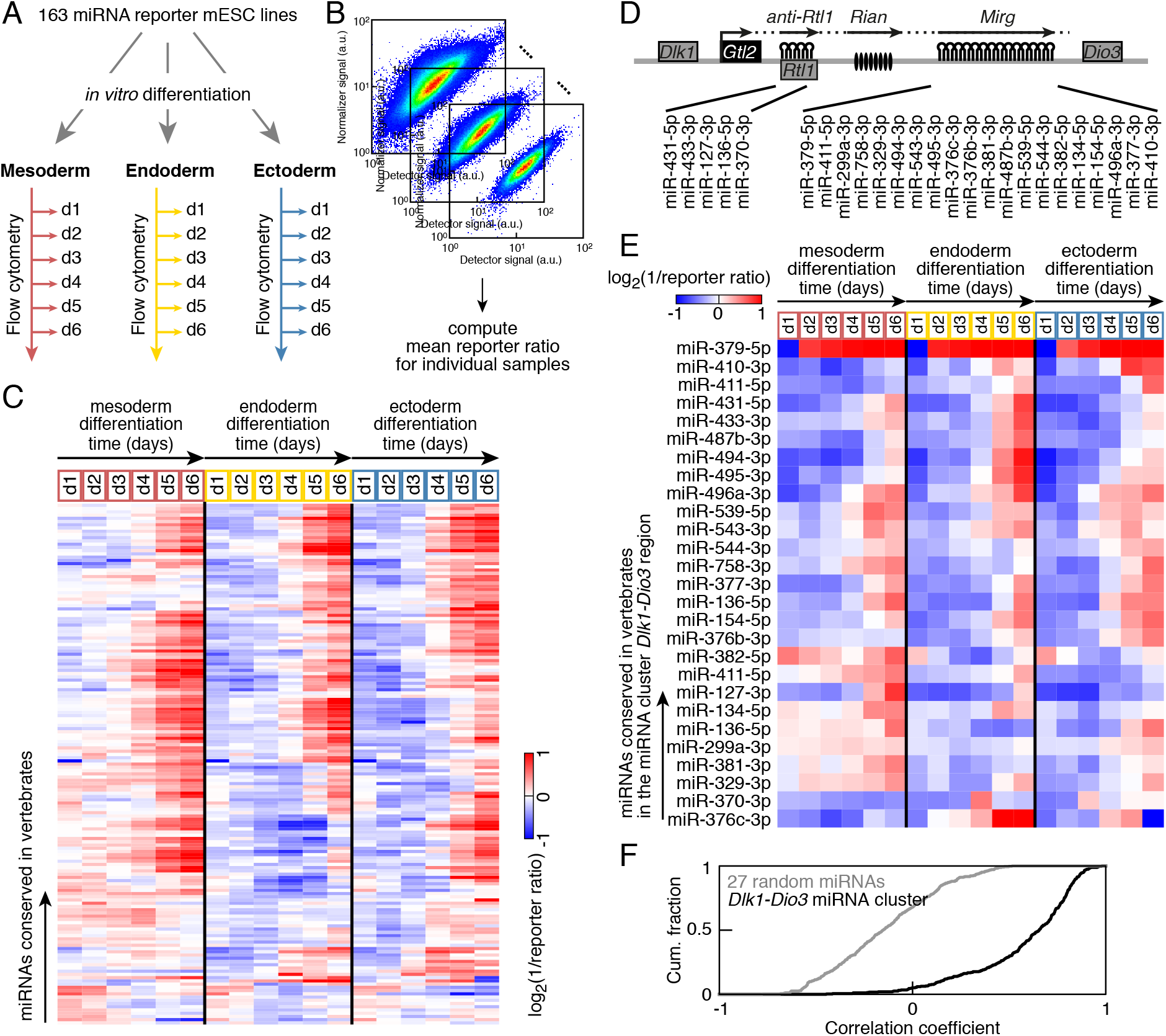
Systematic measurement of miRNA activity during mESC differentiation towards the three germ layers. A Scheme to assess miRNA activity and expression during mESC differentiation towards the three germ layers. B The mean reporter ratio is used as a proxy for miRNA activity in an entire population. C Reporter ratio of miRNAs conserved in vertebrates during mESC differentiation towards mesoderm, endoderm and ectoderm as measured by flow cytometry. D miRNA cluster in the *Dlk1-Dio3* genomic region. The ones conserved in vertebrates are highlighted. E Reporter ratio of miRNAs in the *Dlk1-Dio3* genomic region and conserved in vertebrates during mESC differentiation towards mesoderm, endoderm and ectoderm as measured by flow cytometry. F Cumulative distribution of the correlation coefficient between the reporter ratio of miRNAs in the *Dlk1-Dio3* genomic region and conserved in vertebrates or a random set of 27 miRNAs. p= 10^−91^ for comparison with a random set, Kolmogorov-Smirnov test.

To show the reliability of our activity measurements, we relied on the coregulation of miRNAs belonging to the same genomic cluster which are processed from the same primary transcript. The *Dlk1-Dio3* imprinted region contains the largest miRNA cluster in the mouse genome (Labialle *et al*, 2014). 27 miRNAs of that cluster are conserved in vertebrates and were thus present in our collection of reporter lines (Fig 2D). We found that the activity of the vast majority was gradually upregulated during differentiation to the three germ layers (Fig 2E). More importantly, the changes in activity of miRNAs belonging to the *Dlk1-Dio3* miRNA cluster were highly correlated (Fig 2F), while a set of miRNAs chosen at random from the 163 conserved miRNAs did not show such a correlation (p=10^−91^, Kolmogorov-Smirnov test). This demonstrated both the reliability of our miRNA activity measurements across different stable cell lines and that the majority of miRNAs conserved in vertebrates have a measurable biological activity.

The response curve of our reporter shows that the combination of miRNA expression levels and miRNA reporter ratio allows to determine the target affinity K_D_ for specific miRNAs (Fig 1D). miRNA profiles were measured by deep sequencing every day during differentiation to ectoderm, endoderm and mesoderm. Similarly, the 163 reporter lines were differentiated and the reporter ratio distribution was assessed by flow cytometry, giving us 18 measurements per cell line. We computed K_D_ for individual miRNAs by adjusting the 18 values of mean reporter ratios from miRNA reporter lines with a Hill function depending on the expression levels measured by miRNA-Seq (Fig 3A). Target affinity could be measured for 96 miRNAs with variation in expression during mESC differentiation to the three germ layers (Table 1). K_D_ between miRNAs spanned several orders of magnitude, with an overall positive correlation with median expression levels across mouse tissues (Pearson’s r=0.40, p=4.10^−5^, Fig 3B). In other words, highly expressed miRNAs were on average less potent than weakly expressed miRNAs.

**Table 1.** Target affinity (K_D_) of miRNAs conserved in vertebrates. n.m.: not measurable under our experimental conditions. n.e.: miRNA not expressed under our experimental conditions. RPM: reads per million mapped reads.

| miRNA | K<sub>D</sub> (RPM) | miRNA | K<sub>D</sub> (RPM) | miRNA | K<sub>D</sub> (RPM) | miRNA | K<sub>D</sub> (RPM) |
| --- | --- | --- | --- | --- | --- | --- | --- |
| let-7a-5p | 43000 | miR-137-3p | 60 | miR-216a-5p | 300 | miR-383-5p | 80 |
| miR-1a-3p | 5800 | miR-138-5p | 2000 | miR-216b-3p | n.e. | miR-384-3p | 200 |
| miR-7a-5p | 41000 | miR-139-5p | n.m. | miR-217-3p | n.e. | miR-410-3p | 4400 |
| miR-9-5p | 29000 | miR-140-3p | n.m. | miR-218-5p | 31000 | miR-411-3p | 1900 |
| miR-10b-5p | 190 | miR-141-3p | 450 | miR-219-5p | 3000 | miR-411-5p | 4500 |
| miR-15a-5p | 2100 | miR-142-3p | 200 | miR-221-3p | 1000 | miR-421-3p | 800 |
| miR-17-5p | n.m. | miR-142-5p | 300 | miR-223-3p | 80 | miR-425-5p | n.m. |
| miR-18a-5p | 370 | miR-143-3p | 29000 | miR-224-5p | 3500 | miR-431-5p | n.m. |
| miR-19a-3p | 11000 | miR-144-3p | 130 | miR-290-5p | n.m. | miR-433-3p | 760 |
| miR-21a-5p | 140000 | miR-145a-5p | 460 | miR-292-5p | 470000 | miR-450a-5p | 12000 |
| miR-22-3p | 7100 | miR-146b-5p | n.m. | miR-293-3p | 20000 | miR-451a | 50 |
| miR-23a-3p | 1200 | miR-147-3p | n.e. | miR-294-3p | 98000 | miR-455-5p | 1600 |
| miR-24-3p | 2900 | miR-148a-3p | 110000 | miR-295-3p | 18000 | miR-486-5p | 110 |
| miR-26a-5p | 26000 | miR-149-5p | 450 | miR-296-3p | n.m. | miR-487b-3p | 2600 |
| miR-27b-3p | 11000 | miR-150-5p | 1300 | miR-299a-3p | n.m. | miR-488-3p | 170 |
| miR-29b-3p | n.m. | miR-153-3p | 30 | miR-301a-3p | 230 | miR-488-5p | 180 |
| miR-30c-5p | 45000 | miR-154-3p | n.m. | miR-302b-3p | n.m. | miR-490-3p | 100 |
| miR-31-5p | 910 | miR-155-5p | 430 | miR-302c-3p | 3000 | miR-491-5p | n.e. |
| miR-33-5p | 40 | miR-181a-5p | 5400 | miR-320-3p | n.m. | miR-494-3p | n.m. |
| miR-34c-5p | 4000 | miR-182-5p | n.m. | miR-324-5p | 60 | miR-495-3p | 15000 |
| miR-92a-3p | 28000 | miR-183-5p | n.m. | miR-328-3p | 1000 | miR-496a-3p | n.e. |
| miR-93-5p | n.m. | miR-184-3p | 80 | miR-329-5p | n.m. | miR-499-5p | 100 |
| miR-96-5p | 12000 | miR-186-5p | 1500 | miR-330-5p | 90 | miR-503-5p | 440 |
| miR-99b-5p | 15000 | miR-187-3p | 380 | miR-335-5p | 42000 | miR-504-5p | 25 |
| miR-101b-3p | 6000 | miR-190a-5p | 550 | miR-338-3p | 25 | miR-505-3p | 30 |
| miR-103-3p | 11000 | miR-191-5p | n.m. | miR-339-5p | 600 | miR-539-5p | n.m. |
| miR-122-5p | 12000 | miR-192-5p | 240 | miR-340-5p | 2100 | miR-542-3p | 4300 |
| miR-124-3p | 400 | miR-193b-3p | n.m. | miR-342-3p | 1600 | miR-543-3p | 26000 |
| miR-124-5p | 1400 | miR-194-5p | 950 | miR-346-5p | n.m. | miR-544-3p | 15 |
| miR-125a-3p | 1100 | miR-196a-5p | 1800 | miR-361-5p | 700 | miR-551b-3p | n.e. |
| miR-125a-5p | 370000 | miR-199a-5p | 6500 | miR-365-3p | 80 | miR-590-3p | n.e. |
| miR-126-3p | 2000 | miR-200a-3p | n.m. | miR-370-3p | 5000 | miR-592-3p | n.e. |
| miR-127-3p | 250000 | miR-200b-3p | 16000 | miR-374b-5p | n.m. | miR-615-3p | 2700 |
| miR-128-3p | 7400 | miR-202-3p | n.e. | miR-375-3p | 8100 | miR-653-3p | n.e. |
| miR-129-5p | 8400 | miR-203-3p | n.m. | miR-376b-3p | 270000 | miR-708-5p | n.m. |
| miR-132-3p | 90 | miR-205-5p | n.m. | miR-376c-3p | 940 | miR-758-5p | n.m. |
| miR-133b-3p | 690 | miR-208b-3p | 70 | miR-377-3p | 440 | miR-873b | n.e. |
| miR-134-5p | 25000 | miR-210-3p | n.m. | miR-378c | 2900 | miR-874-3p | n.m. |
| miR-135b-5p | n.m. | miR-211-5p | n.m. | miR-379-5p | 40000 | miR-875-5p | n.e. |
| miR-136-3p | 2000 | miR-212-5p | n.m. | miR-381-3p | n.m. | miR-876-5p | n.e. |
| miR-136-5p | 900 | miR-214-3p | 650 | miR-382-5p | n.m. |  |  |

**Figure 3.**
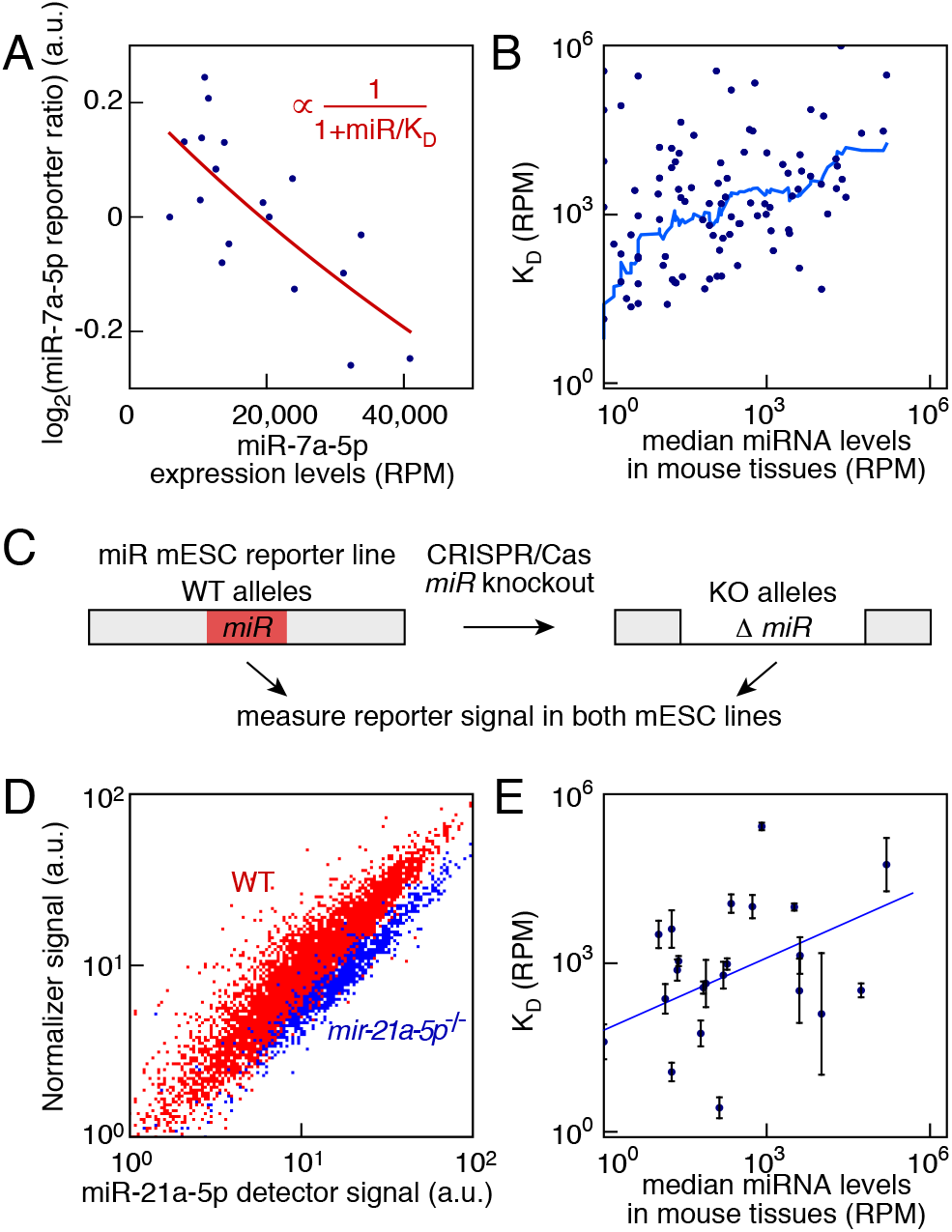
Systematic measurement of target affinity of miRNAs conserved in vertebrates. A Adjustment of reporter ratio as a function of miR-7a-5p levels (blue dots: experimental data measured by miRNA-Seq and flow cytometry) with Hill’s equation (red line). K_D_ is the target affinity. B Comparison between miRNA target affinity K_D_ and miRNA levels in adult tissues (blue line: sliding median average). C Strategy to measure the activity of a miRNA by CRISPR/Cas9-mediated knockout in mESC reporter lines. D miR-21a-5p reporter signal in *miR-21a-5p*^-/-^ mESCs (blue) and in the parental wild type (WT) unedited reporter line (red). E Comparison between miRNA target affinity K_D_ measured by CRISPR/Cas knockout and miRNA levels in adult tissues (Pearson’s r=0.46, p=0.02, error bars represent the confidence interval of one standard deviation on the expression level measurement).

The previous measurements of reporter affinities relied on changes in miRNA expression levels and therefore cannot be applied to constitutively expressed miRNAs. This can be circumvented by controlled artificial changes in expression levels, namely removal by CRISPR/Cas9 of one or both copies of the miRNA to be assessed (Fig 3C). 20 miRNAs were deleted in their corresponding reporter line (Fig S4A–U). We measured the relief in miRNA repression by comparing the reporter ratio in the knockout line and its parental line under pluripotency conditions. As a control, we deleted a conserved miRNA that was not detected in our miRNA-Seq dataset, miR-491-5p. The reporter ratio remained unchanged in *miR-491-5p*^-/-^ mESCs as expected (Fig S4V). Taking the miR-21a-5p reporter line as an example, there was a clear increase in detector signal at the population level in *miR-21a-5p*^-/-^ mESCs compared to the wild type unedited line (Fig 3D). The target affinity was then directly computed using the miRNA expression levels in mESCs (Table 1). As for the previous measurements, affinities spanned several orders of magnitude (Fig S4W). There was a significant correlation between the affinity and median expression levels of that particular miRNA in mouse tissues (Fig 3E), i.e. miRNAs that tend to be more abundant bind more weakly to their targets.

Our large dataset allowed us to ask if the affinity depended on the seed sequence. We predicted the Turner binding energy of seed sequence pairing (Mathews *et al*, 2004). We found no correlation between K_D_ and Δ*G* (Fig S5A). Similarly, there was no correlation between K_D_ and Δ*G* computed for the pairing of nucleotides 2-4 (Fig S5B), nucleotides with which the AGO2-miRNA complex initiates binding (Schirle *et al*, 2014). This is in line with the observation that AGO2 reconfigures the binding energy landscape of its miRNA guide (Salomon *et al*, 2015).

We then investigated whether differences in target affinity K_D_ between miRNAs would have an influence on endogenous targets. Argonaute-CLIP measures the complement of miRNA-bound mRNAs and therefore Ago-CLIP datasets quantify the biologically active miRNAs. Analyzing published data by Bosson et al. (Bosson *et al*, 2014), we found that the total number of Ago-bound 7-mer and 8-mer target sites did not correlate with the expression levels of the corresponding seed family (Pearson’s r=0.09, p=0.52, Fig 4A). For example, the miR-294 and miR-15 families bound the same number of sites despite a 37-fold difference in the expression levels of their respective family members. Even more dramatically, the three miRNA families of miR-142-3p, miR-379 and miR-293 had expression levels spanning more than 2000-fold and yet bound a similar number of targets. We wondered if differences in K_D_ between miRNAs could account for the apparent discrepancy between expression levels and the size of the pool of miRNAs engaged in complexes with Ago and their mRNA targets. Out of the 50 miRNA families represented in the iCLIP dataset, we possessed an affinity value for 24. Like the larger set of 50, expression levels of these miRNAs were not correlated with the number of Ago-bound target sites (Pearson’s r=-0.11, p=0.60). In fact, the target affinity K_D_ should be a determining factor to assess the number of bound miRNAs. We therefore scaled the expression levels of a given miRNA family sharing the same seed by its measured K_D_, yielding “effective” expression levels. Remarkably, the number of bound target sites measured by iCLIP correlated with the effective expression levels for these 24 miRNA families (Pearson’s r=0.49, p=0.016, Fig 4B).

**Figure 4.**
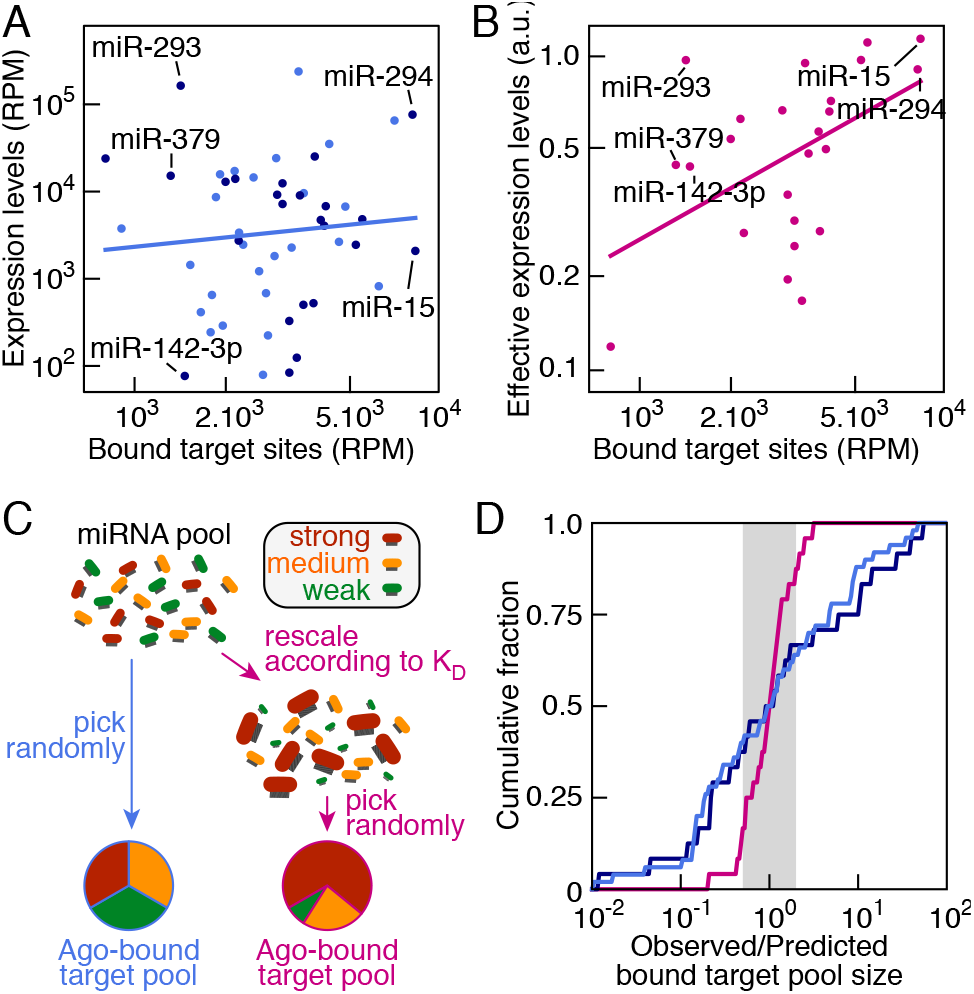
Differences in target affinities account for the distribution of number of bound target sites. A Comparison between the number of bound target sites from a published iCLIP dataset (Bosson *et al*, 2014) and expression levels of the corresponding seed families (dark blue: seeds of conserved miRNAs with measured affinities). B Comparison between the number of bound target sites from a published iCLIP dataset (Bosson *et al*, 2014) and expression levels of the corresponding seed families scaled by their target affinity K_D_ (effective expression). C Models to explain the relative sizes of the Ago-bound target pools. In the first case, miRNAs are chosen randomly according to expression levels. In the second case, miRNA expression levels are first rescaled by the target affinity K_D_ of the respective miRNAs and miRNAs are picked randomly according to their effective expression. D Cumulative fraction of the ratio of the observed number of target sites to the predicted one which target sites are chosen randomly according to miRNA expression levels (light blue: 50 top miRNAs in iCLIP dataset, dark blue: 24 miRNAs with measured K_D_) or to their levels scaled by their target affinity K_D_ (effective expression) (magenta) (p=3.10^−5^, Levene test). Gray shading indicates a 2-fold prediction error.

In order to predict the fraction of bound target sites for individual miRNAs, we constructed two mathematical models that did not possess any adjustable parameters. miRNAs engaging in target repression were selected randomly according either to their absolute expression levels or to their effective concentration (Fig 4C). The model with the effective concentration did significantly better than the random sampling one with a 4.6-fold decrease in the variance (p=3.10^−5^, Levene test). It approximated very well the observed number of bound target sites (Fig 4D): the predicted value was within a 45% error of the actual observed value for 50% of miRNAs and within a 2.44-fold error for 90% of miRNAs.

Despite the simplicity of our mathematical model relying on effective miRNA concentrations, it allowed us to make predictions based on the observation that miRNAs had a higher affinity 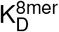 for 8-mer sites compared to the one for 7-mer sites 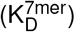 (Nielsen *et al*, 2007; Friedman *et al*, 2009; McGeary *et al*, 2019). First, the fact that the total number of 7- and 8-mer bound sites was well approximated with a single target affinity necessitated that 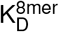 and 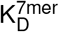 should be proportional to K_D_ for individual miRNAs. When we discriminated between 7- and 8-mer bound sites, we indeed found their numbers to be well correlated with the effective miRNA concentration (Fig S5C–H). This implied that the scaling factor between 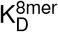 and 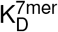 should be comparable for all miRNAs. This prediction was borne out experimentally (McGeary *et al*, 2019). The second was that the numbers of 8-mer and 7-mer bound sites for individual miRNA families should be correlated. It was indeed the case (Pearson’s r=0.75, p=3.10^−10^, Fig S5I). Thus, the affinity measurements made with a synthetic reporter were directly relevant for actual endogenous targets and could explain the relative fraction of bound target sites captured by Ago-iCLIP.

We could measure the *in vivo* target affinity K_D_ for 115 miRNAs out of 163 conserved miRNAs by relying on single-cell fluorescent mESCs reporter lines and physiological changes of the endogenous miRNA levels. 13 miRNAs were not expressed in our experimental conditions while 35 had either too low expression levels to enable a reliable measurement or too many copies in the mouse genome making their deletions unfeasible. K_D_ of individual miRNAs spanned several orders of magnitude. Whereas studies have highlighted the large differences in activity between miRNAs (Mullokandov *et al*, 2012), direct comparisons of miRNA affinities remain rare (McGeary *et al*, 2019). *In vitro* single molecule studies found that let-7 was 2.9 more potent than miR-21 (Salomon *et al*, 2015). We found similarly that let-7 was more potent *in vivo* than miR-21 by a factor of 3.25 (Table 1), representing only a 10% difference between the two estimations relying on different experimental assays.

Remarkably, differences in K_D_ accounted for the relative number of bound target sites for individual miRNA families in a published Ago-CLIP dataset (Bosson *et al*, 2014). For example, miR-15/16 target sites were overrepresented in iCLIP-data compared to the ones of miR-294 (Bosson *et al*, 2014) and indeed this overrepresentation could be explained by the much lower K_D_ of miR-15/16 compared to the one of miR-294 (Table 1). The composition of Ago-bound endogenous sites was quantitatively captured by a simple mathematical model in which miRNAs were chosen randomly according to their effective concentration (defined as their expression levels scaled by their respective K_D_). The model predictions matched very well the independent experimental data without any adjustable parameter. Because this effective concentration ignored differences in relative affinities between different target sites for a given miRNA, a direct prediction of the model was that the numbers of bound 7-mer and 8-mer target sites for a given miRNA family should be proportional. These quantities were indeed correlated. The excellent match between reporter-based K_D_ measurements and CLIP target abundance validates the reliability of our dataset. Moreover, our model suggests that it would be possible to extrapolate relative target affinities for individual miRNAs from the difference in coverage observed by Ago-CLIP.

Contrary to what is commonly reported in other studies using miRNA sensors (Mullokandov *et al*, 2012; Gam *et al*, 2018), the vast majority of the miRNAs conserved in vertebrates had a measurable biological activity. There is a simple explanation to this apparent conundrum: indeed a specific miRNA needs expression levels of the order of K_D_ or greater to have any significant chance to be bound to an accessible site. In fact, the majority of miRNAs are expressed at levels below that threshold in differentiated tissues (Fig 3B) and therefore have limited biological activity. Thus, absolute expression levels are not necessarily predictive of effective target suppression. miR-142 constitutes a prime example of such a miRNA: despite its low expression levels in mESCs, it has a major functional role in mESCs (Sladitschek and Neveu, 2015) because of the small values of the K_D_ of its two mature forms. It follows that the “functional” miRNome will be cell type-dependent, according to the miRNA expression levels in relation to their respective K_D_. This explains previous findings that the very same miRNA can have a biological activity in one cellular context but not another (Mullokandov *et al*, 2012; Bosson *et al*, 2014). The widespread changes of activity we observed during germ layer specification can be linked to the fact that mESCs need a functional miRNA machinery to acquire a differentiated fate. Indeed, both *Dicer*^−*/*−^ mESCs (Kanellopoulou *et al*, 2005; Murchison *et al*, 2005) and *Dgcr8*^−*/*−^ mESCs (Wang *et al*, 2007) fail to differentiate.

Previous assessments of miRNA activity were done in differentiated cell lines (Mullokandov *et al*, 2012; Gam *et al*, 2018) and therefore could not entangle the relative contributions of the expression levels or of the target affinity of a given miRNA to its actual measured activity. As shown in this study, quantitative perturbations of the miRNA levels are needed to decouple the two. Another possible masking factor is the use of multiple miRNA binding sites in reporters (Mullokandov *et al*, 2012; Gam *et al*, 2018), which might render the reporter response ultra-sensitive to the miRNA levels.

Finally, the large span of affinities between individual miRNAs might be of direct relevance for determining the pool of actual mRNA targets. Indeed, 3’-UTRs contain a large number of predicted conserved miRNA binding sites (Grimson *et al*, 2007). Provided that the sites are accessible, one can envision competition between miRNAs for binding of a common target: the combination of relative affinities and expression levels of candidate miRNAs will determine which miRNA will repress that target.

In conclusion, we determined *in vivo* the target affinity K_D_ of miRNAs conserved in vertebrates. K_D_ values varied widely between individual miRNAs and are the yardstick to which expression levels should be compared in order to assess the extent of miRNA repression. Indeed, differences in K_D_ between miRNAs captured the relative size of their respective Argonaute-bound endogenous target pools. Moreover, our collection of 163 stable miRNA reporter mESC lines covering the conserved miRNAs in vertebrates will be a valuable resource for the field.

## Materials and Methods

### miRNA reporter constructs

A single binding site fully complementary to the miRNA to be assessed was cloned downstream of the H2B-Citrine signal in a reporter construct relying on a bidirectional CAG promoter with four CMV enhancers or on a bidirectional PGK promoter (Sladitschek and Neveu, 2016). Wild type and *gp130* 3’-UTRs were as described (Sladitschek and Neveu, 2015) and cloned downstream of the coding sequences of H2B-Citrine and H2B-mCherry.

### mESC maintenance

mESCs were R1 Nagy *et al* (1993) (a kind gift by the EMBL Heidelberg Transgenic Services), E14TG2a (ATCC CRL-1821) or AB2.2 (ATCC SCRC-1023). mESCs were maintained in “LIF+serum” as described previously (Sladitschek and Neveu, 2015).

### Establishment of stable reporter mESC lines

Plasmids were linearized and transfected using Fugene HD (Promega) according to the manufacturer’s protocol. After antibiotic selection, single colonies were expanded and screened for expression of the transgene.

### Generation of miRNA knockout mESCs

Guide RNA inserts targeting miR-9-5p (5’-ATCTAGCTGTATGAGTGGTG), miR-10b-5p (5’-TATAGACAACGTTACAACCT), miR-18a-5p (5’-GCTGCGTGCTTTTTGTTCTA), miR-21a-5p (5’-ATGTTGACTGTTGAATCTCA), miR-92a-3p (5’-TATGGTATTGCACTTGTCC), miR-92b-3p (5’-GGACGAGTGCAATATTGGCG), miR-96-5p (5’-GCAGCCCGCTTTTCCCATAT), miR-99b-5p (5’-GCGATGGTGAGCCCCCGACA), miR-127-3p (5’-TCCGTCTGAGCTTGGCTGGT), miR-136-3p (5’-TATCATGTCGTCGGTTGGAA), miR-181a-5p (5’-CAGTG-AACATTCAACGCTGT), miR-191-5p (5’-AATGGCTGGACAGCGGGCAA), miR-379-5p (5’-GGTGCCCTCC-GAGGATGGAT), miR-433-3p (5’-CGTACTTCTCCCCGGGCATT), miR-451a-3p (5’-CTCTTCTTGGCACAGTTTGT), miR-486a-5p (5’-AAGATCTTCGTTGCGTAGCC), miR-491-5p (5’-GATATGACTTCAACTCAGTC), miR-495-3p (5’-CAACTTCTTTTCAGGTACCA), miR-499-5p (5’-GCCCTACCCAGGCAGCATGG), miR-542-3p (5’-GCAGGGATCTCAGACGTCTC) and miR-615-3p (5’-CCGGCTCGGCCAGTGCTCGG) were designed and cloned in pX330-U6-Chimeric-BB-CBh-hSpCas9 following Hsu *et al* (2013). The Cas9 plasmids were transfected in their respective miRNA reporter mESC line. Successfully edited clones corresponding to *miR-9-5p*^-/-^, *miR-10b-5p*^-/-^, *miR-18a-5p*^-/-^, *miR-21a-5p*^-/-^, *miR-92a-3p*^-/-^, *miR-92b-3p*^-/-^, *miR-96-5p*^-/-^, *miR-99b-5p*^-/-^, *miR-127-3p*^-/-^, *miR-136-3p*^-/-^, *miR-181a-5p*^-/-^, *miR-191-5p*^-/-^, *miR-379-5p*^-/-^, *miR-433-3p*^-/-^, *miR-451a-3p*^-/-^, *miR-486a-5p*^+/-^, *miR-491-5p*^-/-^, *miR-495-3p*^-/-^, *miR-499-5p*^-/-^, *miR-542-3p*^-/-^ and *miR-615-3p*^-/-^ mESCs were validated by genomic PCR.

### mESC differentiation

mESCs were differentiated towards an endodermal progenitor fate (Borowiak *et al*, 2009). Briefly, mESCs were seeded at a density of 2500 cells per cm^2^ onto 0.1% gelatin coated dishes one day prior to the start of the differentiation procedure. The following day, cells were rinsed in D-PBS and switched to endodermal differentiation medium (Advanced RPMI 1640 (ThermoFisher), 1 *µ*M IDE-1 (Tocris), 0.2% (v/v) fetal calf serum (Millipore), 2 mM L-glutamine (Sigma)). Samples collected 24 h after switching to the differentiation regime are referred to as “day 1” differentiation samples. Medium was replaced every day.

mESCs were differentiated towards a mesodermal progenitor fate (Torres *et al*, 2012). Briefly, mESCs were seeded at a density of 2500 cells per cm^2^ onto 0.1% gelatin coated dishes one day prior to the start of the differentiation procedure. The following day, cells were rinsed in D-PBS and switched to mesodermal differentiation medium (Glasgow’s MEM (ThermoFisher), 10% (v/v) KnockOut Serum Replacement (ThermoFisher), 0.1 mM 2-mercaptoethanol (Invitrogen), 1x non-essential amino acids (Gibco) and 1 mM sodium pyruvate (Gibco)). Medium was replaced every day.

mESCs were differentiated towards a neuroectodermal progenitor fate (Ying *et al*, 2003). Briefly, mESCs were seeded at a density of 7500 cells per cm^2^ onto 0.1% gelatin coated dishes one day prior to the start of the differentiation procedure. The following day, cells were washed in D-PBS and switched to N2B27 medium (N2B27 medium was prepared from a 1:1 mixture of DMEM/F12 (without HEPES, with L-glutamine) and neurobasal medium with 0.5x B-27 (with vitamin A) and 0.5x N-2 supplements, 0.25 mM L-glutamine, 0.1 mM 2-mercaptoethanol (all Invitrogen), 10 *µ*g/ml BSA fraction V and 10 *µ*g/ml human recombinant insulin (both Sigma)). *all*-*trans*-Retinoic acid (Sigma) was added at 1 *µ*M to the differentiation medium 24 h after the start of the differentiation procedure. Medium was replaced every other day.

### Flow cytometry

Cells were trypsinized and dissociated to single-cell suspension. Cells were pelleted at 1000g for 1 min, resuspended in D-PBS and strained through a 40 *µ*m filter. Cells were analyzed on an LSRFortessa flow cytometer (BD BioSciences).

### Analysis of flow cytometry data

Flow cytometry data was gated using the forward and side scatter signal to remove debris using FlowJo software and further analyzed in Python. The logarithm of the miRNA reporter ratio was computed from the detector and normalizer signals. The mean of the log-transformed reporter ratio was used to summarize the miRNA activity state in an entire population. In the case of miR-142-3p, miR-142-5p and gp130 3’-UTR reporters, the distribution of the log-transformed reporter ratio was adjusted by a sum of two gaussian distributions corresponding to the two mESC subpopulations in the “high” and “low” miR-142 states. Principal component analysis on mean reporter ratios was carried out as described in Neveu *et al* (2010).

### miRNA-seq library construction

RNA was extracted from cells trypsinized from plates using the MirVana kit (Ambion) following the manufacturer’s instructions. miRNA libraries were prepared using NEBNext Multiplex Small RNA Library Prep Set for Illumina (New England Biolabs) following the manufacturer’s instructions. Libraries were run on Illumina HiSeq 2000 in the 50SE regime. Sequencing results are available on ArrayExpress with accession E-MTAB-4904. Additional miRNA-Seq samples from E-MTAB-2831 (Sladitschek and Neveu, 2015) were used.

### miRNA-seq analysis

After trimming, miRNA reads were matched to miRBase release 19 (Griffiths-Jones *et al*, 2008) mouse sequences allowing no mismatch. Read counts were normalized to account for different sequencing depth. The normalization factor was determined by matching median-filtered log-transformed read counts for two samples to the identity line. Principal component analysis was carried out as described in Neveu *et al* (2010). To assess the impact of miRNAs on their targets, we used matched mRNA-Seq (described in Sladitschek and Neveu (2019)) conducted rom the same total RNA. We kept genes with a maximal expression *>*3 reads per million across samples and at least a 4-fold variation in expression. For a given miRNA, we computed the correlation between its expression levels and the mRNA expression levels of its targets predicted by Targetscan (Grimson *et al*, 2007).

### Computation of miRNA target affinity K_D_

The reporter ratio depends as 1*/*(1 + *M/K*_*D*_) where *M* is the miRNA expression level and *K*_*D*_ the binding affinity (Sladitschek and Neveu, 2015). Read counts were added for all miRNAs belonging to the same seed as these miRNAs will all contribute to the repression of the reporter construct. The mean of log-transformed reporter ratios measured by flow cytometry in single cells in a given sample was computed. Reporter ratios under the 18 different differentiation conditions were adjusted as a function of the corresponding read counts for the matching seed with *α/*(1 + *M/K*_*D*_) with only two free parameters *α* and *K*_*D*_. In the case of miRNA knockouts, the affinity K_D_ was determined by the following formula *M*^+*/*+^*/*(*R*^−*/*−^*/R*^+*/*+^ − 1), where *M*^+*/*+^ is the miRNA expression level in mESCs, *R*^+*/*+^ the reporter ratio in wild type mESCs and *R*^−*/*−^ the reporter ratio in knockout mESCs. In the case of the deletion of one miRNA allele, K_D_ was determined by the following formula (*M*^+*/*+^(1 − 0.5 *R*^+*/*−^*/R*^+*/*+^)*/*(*R*^+*/*−^*/R*^+*/*+^ − 1), where *M*^+*/*+^ is the miRNA expression level in mESCs, *R*^+*/*+^ the reporter ratio in wild type mESCs and *R*^+*/*−^ the reporter ratio in heterozygous mESCs.

### Analysis of iCLIP data from Bosson *et al* (2014)

To compare our miRNA target affinity K_D_ measurements with *in vivo* miRNA-bound target sites, we used iCLIP and miRNA-Seq data in mESCs from Bosson *et al* (2014) with accession number GSE61348. We retained all 8-mer and 7-mer bound target sites with at least one read (a threshold shown to be above background according to Bosson *et al* (2014)) for the top 50 miRNA families. 6-mer sites were discarded as they cannot always be attributed unambiguously to a single seed. The associated published miRNA expression dataset was analyzed. miRNA reads that matched to miRBase release 19 (Griffiths-Jones *et al*, 2008) mouse sequences with no mismatch were kept. Read counts of miRNAs sharing the same seed were added to reflect the total expression levels of the corresponding miRNA family. Out of the 50 seeds represented in the data, we possessed affinity measurements for 24. Expression levels of the miRNA family (in reads per million) were scaled by K_D_ of the corresponding seed. Given that the size of the total target pool varies little across miRNAs and that the total target pool of a given miRNA is more abundant than its corresponding miRNA (Bosson *et al*, 2014), we devised two models to explain the relative size of Ago-bound target pools of individual miRNA families. In the first model, Ago-bound miRNAs are picked at random from the total miRNA pool. In the second model, miRNA concentrations are scaled by their respective affinity K_D_ and then Ago-bound miRNAs are picked at random from this rescaled miRNA pool. The predicted numbers of Ago-bound miRNAs are then compared to their experimentally observed target pool size (Bosson *et al*, 2014). As we considered the relative sizes of target pools, our models did not have any adjustable parameters.

## Data deposition

Sequencing results are available on ArrayExpress with accession E-MTAB-4904.

## Statistical analysis

Statistical tests were computed using the Python SciPy module.

## Acknowledgments

We thank Jérôme Sinniger for excellent support with cell culture. This work was technically supported by the EMBL Flow Cytometry and Genomics Core facilities. The study was funded by EMBL. S.B. was also supported by the EMBL International PhD Programme (EIPP).

## Author contributions

S.B. and H.L.S. carried out experiments and analyzed the data. P.A.N. conceived the study, analyzed the data and wrote the paper.

## References

Bartel DP (2004) MicroRNAs: genomics, biogenesis, mechanism, and function. Cell 116: 281–297

Bartel DP (2009) MicroRNAs: target recognition and regulatory functions. Cell 136: 215–233

Borowiak M, Maehr R, Chen S, Chen AE, Tang W, Fox JL, Schreiber SL, Melton DA (2009) Small molecules efficiently direct endodermal differentiation of mouse and human embryonic stem cells. Cell Stem Cell 4: 348–358

Bosson AD, Zamudio JR, Sharp PA (2014) Endogenous miRNA and target concentrations determine susceptibility to potential ceRNA competition. Mol Cell 56: 347—359

Carthew RW, Sontheimer EJ (2009) Origins and Mechanisms of miRNAs and siRNAs. Cell 136: 642–655

Chandradoss SD, Schirle NT, Szczepaniak M, MacRae IJ, Joo C (2015) A Dynamic Search Process Underlies MicroRNA Targeting. Cell 162: 96—107

Chi SW, Zang JB, Mele A, Darnell RB (2009) Argonaute HITS-CLIP decodes microRNA-mRNA interaction maps. Nature 460: 479—486

Ebert MS, Neilson JR, Sharp PA (2007) MicroRNA sponges: competitive inhibitors of small RNAs in mammalian cells. Nat Methods 4: 721–726

Elkayam E, Kuhn CD, Tocilj A, Haase AD, Greene EM, Hannon GJ, Joshua-Tor L (2012) The structure of human argonaute-2 in complex with miR-20a. Cell 150: 100—110

Fabian MR, Sonenberg N, Filipowicz W (2010) Regulation of mRNA translation and stability by microRNAs. Annu Rev Biochem 79: 351–379

Friedman RC, Farh KKH, Burge CB, Bartel DP (2009) Most mammalian mRNAs are conserved targets of microRNAs. Genome Res 19: 92—105

Gam JJ, Babb J, Weiss R (2018) A mixed antagonistic/synergistic miRNA repression model enables accurate predictions of multi-input miRNA sensor activity. Nat Commun 9: 2430

Griffiths-Jones S, Saini HK, Van Dongen S, Enright AJ (2008) miRBase: tools for microRNA genomics. Nucleic Acids Res 36: D154–8

Grimson A, Farh KKH, Johnston WK, Garrett-Engele P, Lim LP, Bartel DP (2007) MicroRNA targeting specificity in mammals: determinants beyond seed pairing. Mol Cell 27: 91–105

Grimson A, Srivastava M, Fahey B, Woodcroft BJ, Chiang HR, King N, Degnan BM, Rokhsar DS, Bartel DP (2008) Early origins and evolution of microRNAs and Piwi-interacting RNAs in animals. Nature 455: 1193–1197

Guo H, Ingolia NT, Weissman JS, Bartel DP (2010) Mammalian microRNAs predominantly act to decrease target mRNA levels. Nature 466: 835–840

Hafner M, Landthaler M, Burger L, Khorshid M, Hausser J, Berninger P, Rothballer A, Ascano M, Jungkamp AC, Munschauer M, Ulrich A, Wardle GS, Dewell S, Zavolan M, Tuschl T (2010) Transcriptome-wide identification of RNA-binding protein and microRNA target sites by PAR-CLIP. Cell 141: 129–141

Hsu PD, Scott DA, Weinstein JA, Ran FA, Konermann S, Agarwala V, Li Y, Fine EJ, Wu X, Shalem O, Cradick TJ, Marraffini LA, Bao G, Zhang F (2013) DNA targeting specificity of RNA-guided Cas9 nucleases. Nat Biotechnol 31: 827–832

Kanellopoulou C, Muljo SA, Kung AL, Ganesan S, Drapkin R, Jenuwein T, Livingston DM, Rajewsky K (2005) Dicer-deficient mouse embryonic stem cells are defective in differentiation and centromeric silencing. Genes Dev 19: 489–501

Khorshid M, Hausser J, Zavolan M, van Nimwegen E (2013) A biophysical miRNA-mRNA interaction model infers canonical and noncanonical targets. Nat Methods 10: 253–255

Labialle S, Marty V, Bortolin-Cavaillé ML, Hoareau-Osman M, Pradère JP, Valet P, Martin PGP, Cavaillé J (2014) The miR-379/miR-410 cluster at the imprinted Dlk1-Dio3 domain controls neonatal metabolic adaptation. EMBO J 33: 2216—2230

Lewis BP, Burge CB, Bartel DP (2005) Conserved seed pairing, often flanked by adenosines, indicates that thousands of human genes are microRNA targets. Cell 120: 15–20

Lim LP, Lau NC, Garrett-Engele P, Grimson A, Schelter JM, Castle J, Bartel DP, Linsley PS, Johnson JM (2005) Microarray analysis shows that some microRNAs downregulate large numbers of target mRNAs. Nature 433: 769–773

Mathews DH, Disney MD, Childs JL, Schroeder SJ, Zuker M, Turner DH (2004) Incorporating chemical modification constraints into a dynamic programming algorithm for prediction of RNA secondary structure. Proc Natl Acad Sci USA 101: 7287—7292

McGeary SE, Lin KS, Shi CY, Pham T, Bisaria N, Kelley GM, Bartel DP (2019) The biochemical basis of microRNA targeting efficacy. Science : aav1741

Mukherji S, Ebert MS, Zheng GXY, Tsang JS, Sharp PA, van Oudenaarden A (2011) MicroRNAs can generate thresholds in target gene expression. Nat Genet 43: 854–859

Mullokandov G, Baccarini A, Ruzo A, Jayaprakash AD, Tung N, Israelow B, Evans MJ, Sachidanandam R, Brown BD (2012) High-throughput assessment of microRNA activity and function using microRNA sensor and decoy libraries. Nat Methods 9: 840–846

Murchison EP, Partridge JF, Tam OH, Cheloufi S, Hannon GJ (2005) Characterization of Dicer-deficient murine embryonic stem cells. Proc Natl Acad Sci USA 102: 12135–12140

Nagy A, Rossant J, Nagy R, Abramow-Newerly W, Roder JC (1993) Derivation of completely cell culture-derived mice from early-passage embryonic stem cells. Proc Natl Acad Sci USA 90: 8424–8428

Nakanishi K, Weinberg DE, Bartel DP, Patel DJ (2012) Structure of yeast Argonaute with guide RNA. Nature 486: 368—374

Neveu P, Kye MJ, Qi S, Buchholz DE, Clegg DO, Sahin M, Park IH, Kim KS, Daley GQ, Kornblum HI, Shraiman BI, Kosik KS (2010) MicroRNA profiling reveals two distinct p53-related human pluripotent stem cell states. Cell Stem Cell 7: 671–681

Nielsen CB, Shomron N, Sandberg R, Hornstein E, Kitzman J, Burge CB (2007) Determinants of targeting by endogenous and exogenous microRNAs and siRNAs. RNA 13: 1894—1910

Pasquinelli AE, Reinhart BJ, Slack F, Martindale MQ, Kuroda MI, Maller B, Hayward DC, Ball EE, Degnan B, Müller P, Spring J, Srinivasan A, Fishman M, Finnerty J, Corbo J, Levine M, Leahy P, Davidson E, Ruvkun G (2000) Conservation of the sequence and temporal expression of let-7 heterochronic regulatory RNA. Nature 408: 86–89

Reinhart BJ, Slack FJ, Basson M, Pasquinelli AE, Bettinger JC, Rougvie AE, Horvitz HR, Ruvkun G (2000) The 21-nucleotide let-7 RNA regulates developmental timing in Caenorhabditis elegans. Nature 403: 901–906

Salomon WE, Jolly SM, Moore MJ, Zamore PD, Serebrov V (2015) Single-Molecule Imaging Reveals that Argonaute Reshapes the Binding Properties of Its Nucleic Acid Guides. Cell 162: 84—95

Schirle NT, MacRae IJ (2012) The crystal structure of human Argonaute2. Science 336: 1037—1040

Schirle NT, Sheu-Gruttadauria J, MacRae IJ (2014) Structural basis for microRNA targeting. Science 346: 608—613

Sladitschek HL, Neveu PA (2015) The bimodally expressed microRNA miR-142 gates exit from pluripotency. Mol Syst Biol 11: 850

Sladitschek HL, Neveu PA (2016) Bidirectional Promoter Engineering for Single Cell MicroRNA Sensors in Embryonic Stem Cells. PloS one 11: e0155177

Sladitschek HL, Neveu PA (2019) A gene regulatory network controls the balance between mesendoderm and ectoderm at pluripotency exit. Mol Syst Biol 15: e9043

Torres J, Prieto J, Durupt FC, Broad S, Watt FM (2012) Efficient differentiation of embryonic stem cells into mesodermal precursors by BMP, retinoic acid and Notch signalling. PloS one 7: e36405

Viswanathan SR, Daley GQ, Gregory RI (2008) Selective blockade of microRNA processing by Lin28. Science 320: 97–100

Wang Y, Medvid R, Melton C, Jaenisch R, Blelloch R (2007) DGCR8 is essential for microRNA biogenesis and silencing of embryonic stem cell self-renewal. Nat Genet 39: 380–385

Wee LM, Flores-Jasso CF, Flores-Jasso CF, Salomon WE, Zamore PD (2012) Argonaute divides its RNA guide into domains with distinct functions and RNA-binding properties. Cell 151: 1055—1067

Ying QL, Stavridis M, Griffiths D, Li M, Smith A (2003) Conversion of embryonic stem cells into neuroectodermal precursors in adherent monoculture. Nat Biotechnol 21: 183–186

