## Supplemental figures for "Systematic *in vivo* quantification of microRNA affinities"

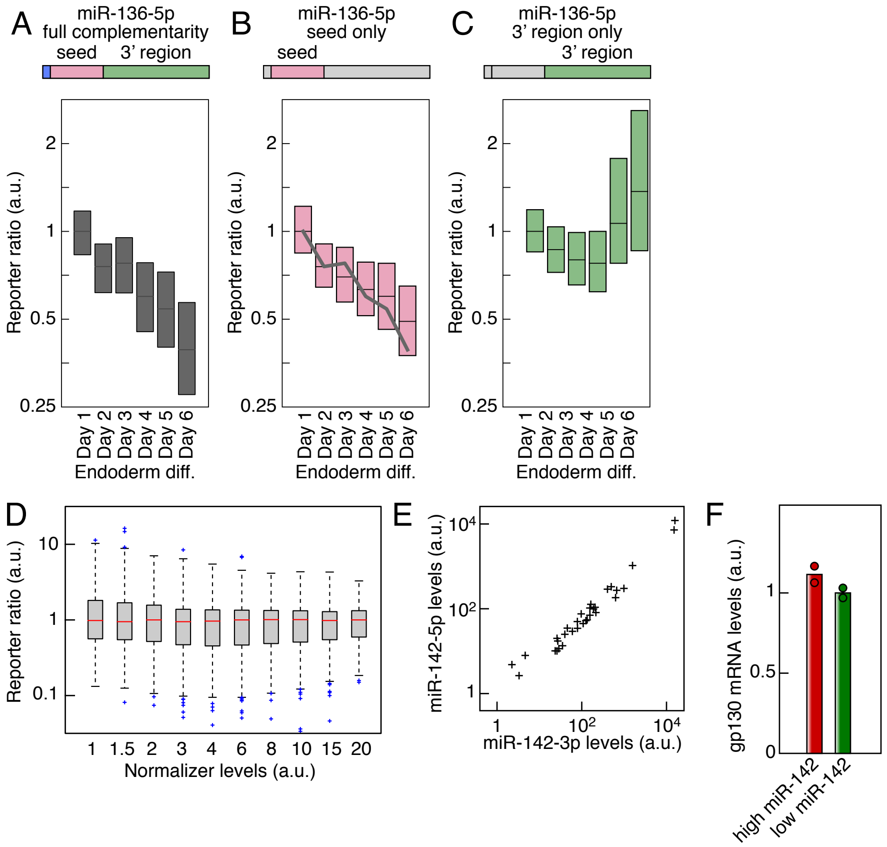


**Figure S1. The miRNA activity reporter is sensitive to the seed region.**

A Box plot of the reporter ratio during mESC differentiation to endoderm for a detector containing a binding site fully complementary to miR-136-5p.

B Box plot of the reporter ratio during mESC differentiation to endoderm for a detector containing a binding site complementary to miR-136-5p seed region (black line: median reporter ratio for the fully complementary reporter).

C Box plot of the reporter ratio during mESC differentiation to endoderm for a detector containing a binding site complementary to the 3' region of miR-136-5p.

D Box and whisker plot of a miR-142-3p activity reporter ratio as a function of the normalizer levels.

E Comparison between miR-142-3p and miR-142-5p expression levels (Pearson's r=0.977, $p=10^{-21}$).

F gp130 mRNA levels in FACS-purified high and low miR-142 populations.

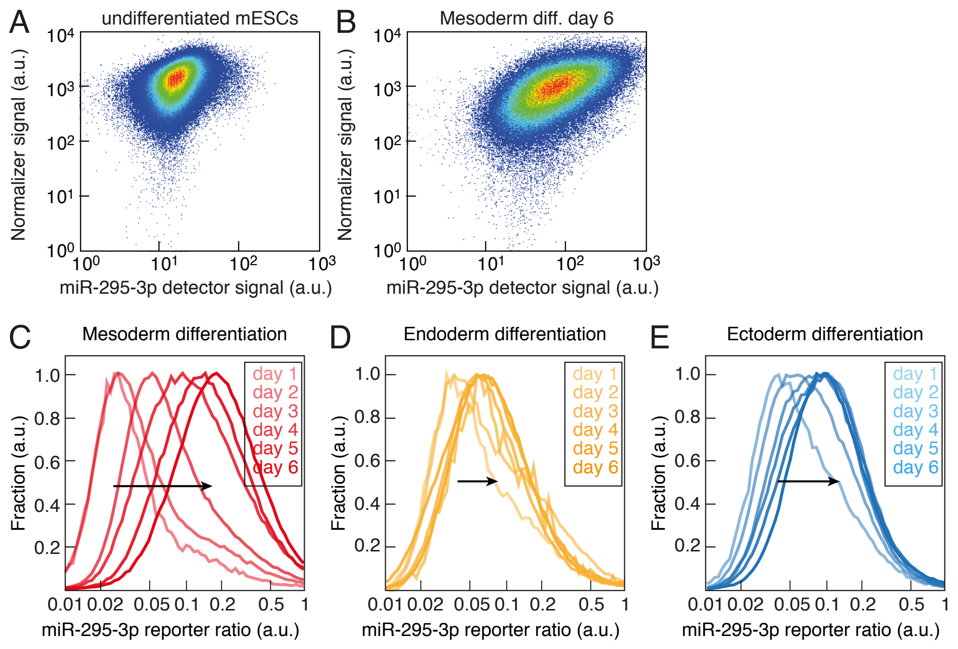


**Figure S2. Changes in miRNA activity can be assessed during mESC differentiation.**

A Detector and normalizer expression in a clonal mESC population stably expressing a miR-295-3p reporter.

B Detector and normalizer expression in mESCs stably expressing a miR-295-5p reporter and differentiated for 6 days to mesoderm.

C Temporal evolution of the distribution of miR-295-3p reporter ratio in mESCs differentiated to mesoderm (line color according to the inset).

D Distribution of miR-295-3p reporter ratio in mESCs differentiated to endoderm (line color according to the inset).

E Distribution of miR-295-3p reporter ratio in mESCs differentiated to ectoderm (line color according to the inset).

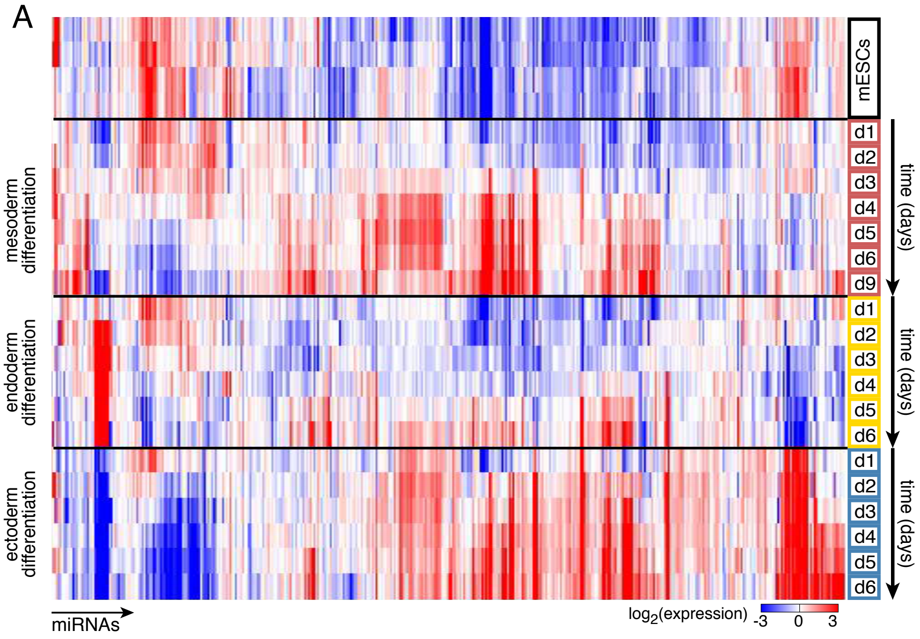


B Number of miRNAs conserved in vertebrates with expression changes

| | mesoderm differentiation | endoderm differentiation | ectoderm differentiation |
|---|---|---|---|
| > 2 fold change | 98 | 59 | 69 |
| > 4 fold change | 33 | 37 | 45 |

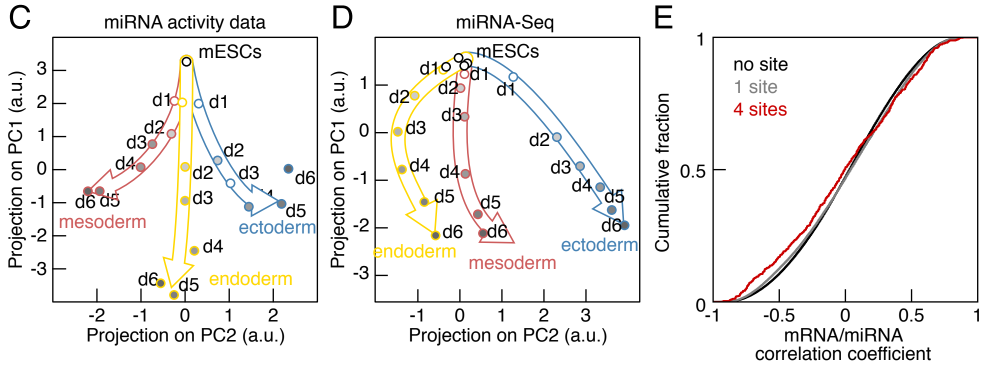


**Figure S3. miRNA expression during mESC differentiation towards the three germ layers.**

A miRNA expression levels in mESCs differentiated towards ectoderm (blue), mesoderm (red) and endoderm (yellow).

B Number of miRNAs conserved in vertebrates with expression changes during differentiation to the three germ layers.

C Principal component analysis of miRNA activity changes as measured with miRNA reporter lines during mESC differentiation (black: mESCs, blue: ectoderm, red: mesoderm, yellow: endoderm).

D Principal component analysis of miRNA expression changes during mESC differentiation (black: mESCs, blue: ectoderm, red: mesoderm, yellow: endoderm).

E Cumulative distribution of the correlation coefficient between the expression profiles of differentially regulated miRNAs and of genes with zero (black), one (gray) or four (red) predicted binding sites for this miRNA in their 3'-UTRs. p=0.6 and $10^{-5}$ for the 0-1 and 0-4 sites comparisons, Kolmogorov-Smirnov test.

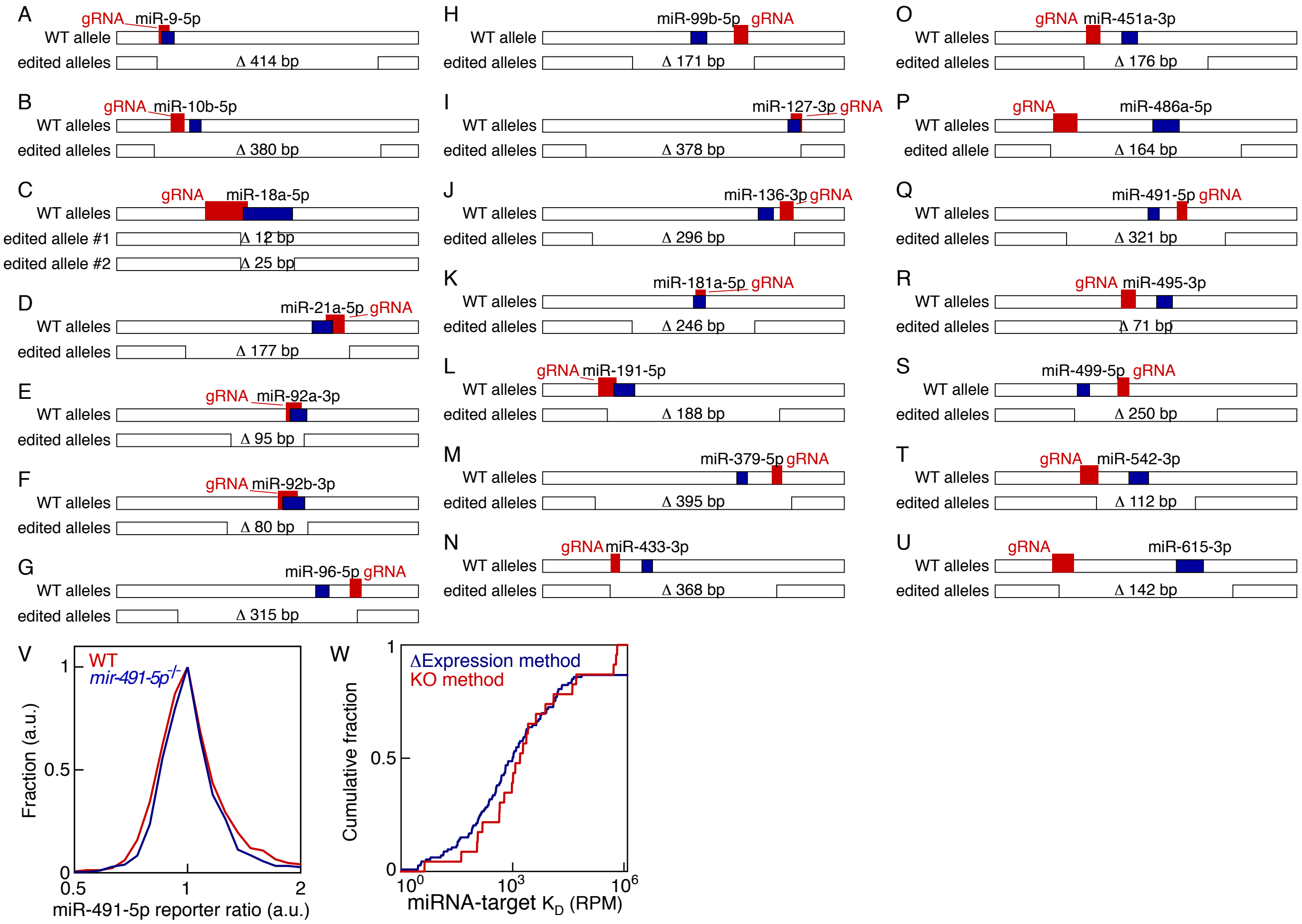


**Figure S4. Generation of miRNA knockout mESC lines.**

A–U Genomic locus and corresponding deletion alleles for miR-9-5p (A), miR-10b-5p (B), miR-18a-5p (C), miR-21a-5p (D), miR-92a-3p (E), miR-92b-3p (F), miR-96-5p (G), miR-99b-5p (H), miR-127-3p (I), miR-136-3p (J), miR-181a-5p (K), miR-191-5p (L), miR-379-5p (M), miR-433-3p (N), miR-451a-3p (O), miR-486a-5p (P), miR-491-5p (Q), miR-495-3p (R), miR-499-5p (S), miR-542-3p (T) and miR-615-3p (U). The position of the miRNA is indicated in blue and the targeting sequence of the guide RNA (gRNA) in red. The size of the deletion is indicated in base pairs (bp).

V miR-491-5p reporter signal in *miR-491-5p*$^{-/-}$ mESCs (blue line) and in the parental wild type (WT) unedited reporter line (red line).

W Cumulative distribution of affinities obtained by mESC differentiation (blue line, ΔExpression Method) or CRISPR/Cas-mediated deletion of miRNAs (red line, KO Method).

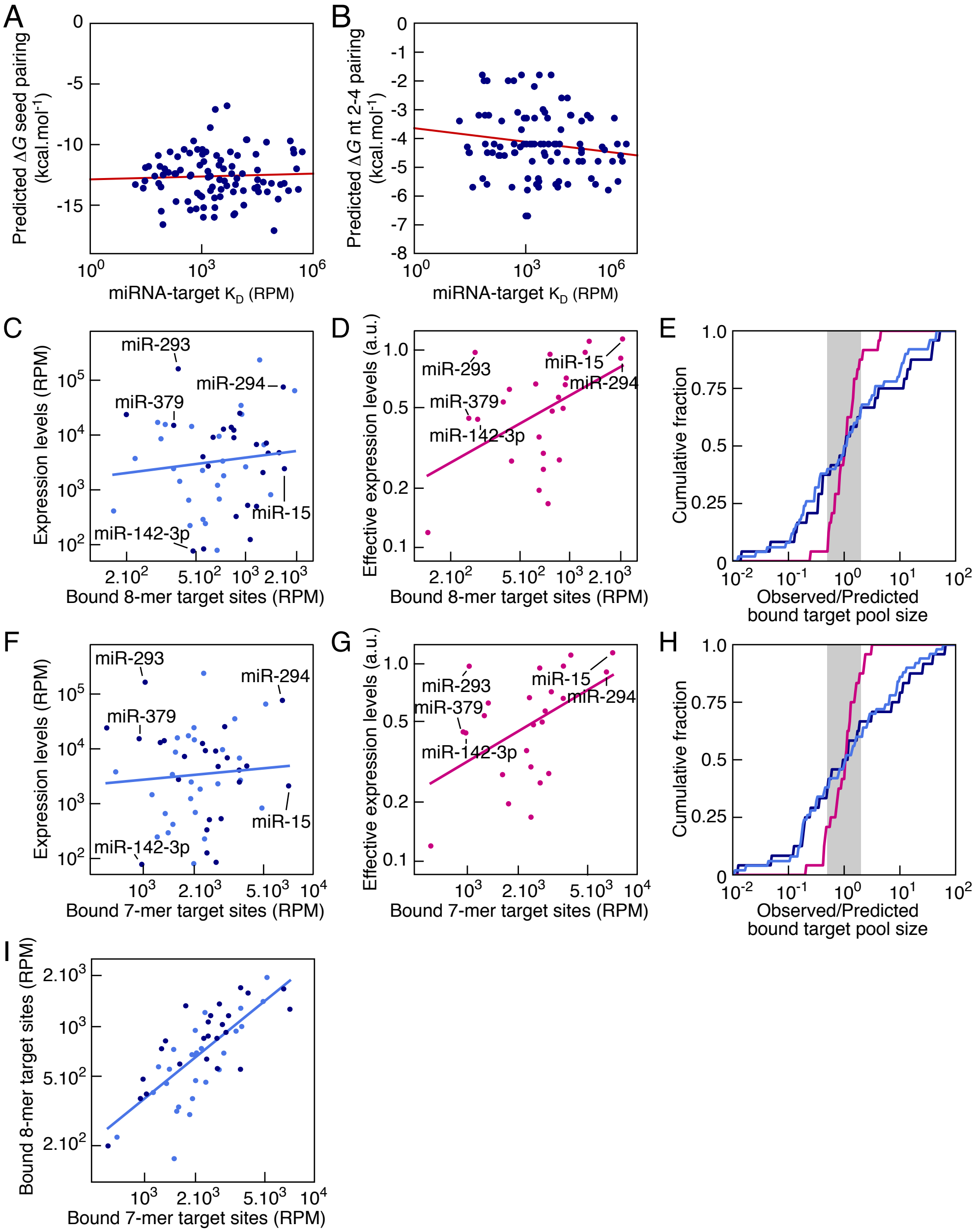


**Figure S5. Differences in target affinities account for the distribution of number of bound target sites.**

A–B Comparison between miRNA target affinities ($K_D$) and predicted $\Delta$G for the pairing of the entire seed (A) or the nucleotides 2-4 (B).

C Comparison between the number of bound 8-mer target sites from a published iCLIP dataset (Bosson *et al*, 2014) and expression levels of the corresponding seed families (dark blue: seeds of conserved miRNAs with measured affinities). Pearson's r=0.11, p=0.42.

D Comparison between the number of bound 8-mer target sites from a published iCLIP dataset (Bosson *et al*, 2014) and expression levels of the corresponding seed families scaled by their target affinity $K_D$ (effective expression). Pearson's r=0.40, p=0.05.

E Cumulative fraction of the ratio of the observed fraction of target 8-mer sites to the predicted one where target sites are chosen randomly according to miRNA expression levels (light blue: 50 top miRNAs in iCLIP dataset, dark blue: 24 miRNAs with $K_D$) or to their levels scaled by their target affinity $K_D$ (effective expression) (magenta) ($p=7.10^{-5}$, Levene test). Gray shading indicates a 2-fold prediction error.

F Comparison between the number of bound 7-mer target sites from a published iCLIP dataset (Bosson *et al*, 2014) and expression levels of the corresponding seed families (dark blue: seeds of conserved miRNAs with measured affinities). Pearson's r=0.08, p=0.58.

G Comparison between the number of bound 7-mer target sites from a published iCLIP dataset (Bosson *et al*, 2014) and expression levels of the corresponding seed families scaled by their target affinity $K_D$ (effective expression). Pearson's r=0.49, p=0.015.

H Cumulative fraction of the ratio of the observed fraction of target 7-mer sites to the predicted one where target sites are chosen randomly according to miRNA expression levels (light blue: 50 top miRNAs in iCLIP dataset, dark blue: 24 miRNAs with $K_D$) or to their levels scaled by their target affinity $K_D$ (effective expression) (magenta) ($p=3.10^{-5}$, Levene test). Gray shading indicates a 2-fold prediction error.

I Comparison between the numbers of bound 8-mer and 7-mer target sites from a published iCLIP dataset (Bosson *et al*, 2014). Pearson's r=0.75, $p=3.10^{-10}$. Dark blue: seeds of conserved miRNAs with measured affinities.
